# Stand Age and Climate Change Effects on Carbon Increments and Stock Dynamics

**DOI:** 10.1101/2024.05.23.595487

**Authors:** Elia Vangi, Daniela Dalmonech, Mauro Morichetti, Elisa Grieco, Francesca Giannetti, Giovanni D’Amico, Mahdi (Andre) Nakhavali, Gherardo Chirici, Alessio Collalti

## Abstract

Carbon assimilation and wood production are influenced by environmental conditions and endogenous factors, such as species auto-ecology, age, and hierarchical position within the forest structure. Disentangling the intricate relationships between those factors is more pressing than ever due to climate change’s pressure. We employed the 3D-CMCC-FEM model to simulate undisturbed forests of different ages under four climate change scenarios (plus one no climate change) from five Earth System Models. In this context, carbon stocks and increment were simulated via total car-bon woody stocks and mean annual increment, which depends mainly on climate trends. We find greater differences among different age cohorts under the same scenario than in different climate scenarios under the same age class. Increasing temperature and changes in precipitation patterns led to a decline in above-ground biomass in spruce stands, espe-cially in the older age classes. On the contrary, the results show that beech forests at DK-Sor will maintain and even increase C-storage rates under most RCP scenarios. Scots pine forests show an intermediate behavior with a stable stock capacity over time and in different scenarios but with decreasing mean volume annual increment. These results confirm current observations worldwide that indicate a stronger climate-related decline in conifers forests than in broadleaves.

## 1. Introduction

Assessing the quantity of CO2 equivalent stored in forest ecosystems is one of the main goals for implementing the new European Forest Strategy for 2030, a key component of the European Green Deal, to achieve greenhouse gas emission neutrality by 2050. Within this framework, European forest strategies have been geared towards forest-based mitigation plans [1, 2], which makes it essential to estimate the carbon sequestration capacity and potential under future climate conditions.

In the near future, Europe and Mediterranean areas will emerge as focal points (’hot spots’) of climate change, characterized by heightened temperatures and environmental impacts [3, 4]. Carbon assimilation and wood production are influenced by environmental conditions (e.g., precipitation, temperature, atmospheric CO2, etc.) and endogenous factors, such as species auto-ecology, age, and hierarchical position within the forest structure. In the past decades, forest ecosystems proved to be a crucial net carbon sinks [5, 6], likely due to the positive fertilization effects of rising atmospheric CO2 and temperature [7]. However, whether this effect will remain positive or be compensated by other limiting factors is still a matter of debate [8, 9, 10]. Some studies suggest that the fertilization effect on carbon storage and biomass production fades with forest aging in temperate forests [11, 12] since these positive effects cannot continue indefinitely, complicating the picture of the forest response to climate changes even further. This is already the case in Europe, where forest aging and increased disturbances are causing the saturation and decline of the forest carbon sink [9]. Unfortunately, there is not yet a clear strategy to increase the mitigation potentials of forests, and the factors involved are manifold and entangled together [11, 13, 14].

The need to disentangle the intricate relationships between those factors is even more pressing under climate change. Our current understanding of how future climate will interact with forests of different age classes is particularly limited, especially since only a few studies have explored the relationship between age and the ecosystem’s carbon balance under changing climate conditions [15].

The climate sensitivity of age cohorts is driven, among all, by different access to environmental resources, such as root depth and, therefore, access to water, as well as height, which affects leaf-level water potential and, thus, stomatal conductance [16]. Rooting depth and height jointly affect the tree’s sensitivity to water scarcity, a key environmental driver of change. Future changes in environmental conditions are expected to impact the age spectrum differently [17, 18, 19].

Since forest age is determined by management practices and 75% of European forests are even-aged [20, 21], it is crucial to grasp and pin down the role of age in the sensitivity of forest carbon stocks to climate change to guide and inform adaptative forest management.

Process-based forest models enable the exploration of climate change impacts on various age cohorts within the same area, a task difficult to achieve through direct field measurements, which would require decades or more. In this regard, this study examines the ability of different forest age classes under the same future climate conditions to sustain high productivity and carbon stock capacity. To achieve this goal, we employed the ‘Three Dimensional - Coupled Model Carbon Cycle - Forest Ecosystem Module’ (3D-CMCC-FEM) [22, 23], simulating undisturbed forests of different cohorts under four climate change scenarios (and including one ‘no climate change’ scenario), from the moderate one (RCP 2.6) up to the most severe one (RCP 8.5) coming from five Earth System Models. In this context, carbon stocks and increment were simulated via total carbon woody stocks (TCWS, i.e., the standing woody biomass in MgC ha^—1^) and the mean annual increment (MAI, in m^3^ ha^—1^year^—1^), which depend mainly on age and long-term processes, such as climate trends.

The primary aim of this research is to explore (i) the direct effects of climate change on the overall carbon storage capacity across various stands, species, and age classes situated in diverse regions of Europe; (ii) elucidate the potential influence of forest age on stand dynamics in adapting to forthcoming climate shifts.

## 2. Materials and Methods

### 2.1. Study sites

The study was conducted in three even-aged, previously managed European forest stands i) the Boreal Scots pine (Pinus sylvestris L.) forest of Hyytiälä, Finland (FI-Hyy); ii) the wet temperate continental Norway spruce (Picea abies (L.) H. Karst) forest of Bílý Krìz in the Czech Republic (CZ-BK1); and iii) the temperate oceanic European beech (Fagus sylvatica L.) forest of Sorø, Denmark (DK-Sor) where the 3D-CMCC-FEM (in different versions) has been already validated in the past [14, 24, 25]. An overview of the main site’s characteristics is presented in tab.1 and 2.

For each site, daily bias-adjusted downscaled climate data from five Earth System Models (i.e., HadGEM2-ES, IPSL-CM5A-LR, MIROC-ESM-CHEM, GFDL-ESM2M, and NorESM1-M) driven by four Representative Concentration Pathways, namely RCP 2.6, 4.5, 6.0, and 8.5 were available [26, 27] (Fig. S1). For more detailed information on the study site characteristics and climate data, see [14, 24, 25, 28]. The chosen sites have been selected due to their long monitoring history and the availability of a wide range of data sources for both carbon fluxes and biometric data for model evaluation, as well as bias-corrected climate scenarios for simulations under climate change scenarios from the ISIMIP-PROFOUND initiatives (https://www.isimip.org/)[25, 28]. In addition, these stands: i) represent the most common European tree species; ii) their current state is the result of the legacy of past forest management; iii) they are mainly mono-specific and therefore represent interesting «living labs» to study the effects of climate change on single-species and their productivity, reducing confounding effects which otherwise make models struggle to predict forest growth and carbon dynamics (e.g., [29, 30]), iv), and they have already been investigated in the context of climate-smart-forestry silvicultural scenarios [14].

### 2.2. The model

The ‘Three Dimensional - Coupled Model Carbon Cycle - Forest Ecosystem Module’ (3D-CMCC-FEM v 5.6 [12, 14, 22, 23, 24, 31] is a biogeochemical, biophysical, process-based, stand-level forest model. The model is built to simulate carbon, nitrogen, and water cycles in forest ecosystems, even including forest dynamics, under scenarios of climate change and disturbances (e.g., forest management) and parameterized at the species level. Photosynthesis is modeled through the biogeochemical model of Farquhar von Caemmerer and Berry [32] implemented for sun and shaded leaves [33] and parametrized as in Bernacchi et al. [34, 35]. Temperature acclimation of leaf photosynthesis to increasing temperature is accounted for following Kattge and Knorr [36].

Autotrophic respiration (RA) is modeled mechanistically by distinguishing the costs of maintaining already existing tissues (RM) and the cost of synthesizing new ones (RG). Maintenance respiration is controlled by the amount of nitrogen (stoichiometrically fixed fraction of live tissues) and temperature. Temperature effects on enzyme kinetics are modeled through a standard Arrhenius relationship but acclimated for temperature as described in Collalti et al. [24]. The net primary productivity (NPP) is the gross primary productivity (GPP) less RA. Not all the annual NPP goes for biomass production since the model considers the Non-structural carbon (NSC) pool, an additional seventh C-pool which includes starch and sugars (undistinguished) used to buffer periods of negative carbon balance (when respiration exceeds assimilation; i.e., RA > GPP). Ultimately, the more trees respire, the more NSC is used to sustain metabolism and NSC pool replenishment, and the less NPP and BP there are (and less carbon is stocked). In the extreme case, when and if all NSCs are depleted because of metabolism without being replenished through current photosynthates, the model predicts stand mortality based on the carbon starvation hypothesis [37, 38].

The phenological and allocation schemes are all described extensively in Collalti et al. [22, 23, 39] and Merganičová et al. [39]. The 3D-CMCC-FEM accounts for the ‘age-effect’ in several ways. ‘60s ecological theories describe [40, 41], and past and growing pieces of evidence suggest, that stabilization and a further slight decline follow an initial step-wise increase in forest productivity.

The causes of such a decline are debated and include a decline in the GPP because of hydraulic limitation []16, 42] or an increase in RA because of increased respiring biomass [18, 19, 43]. The 3D-CMCC-FEM accounts for both by including an age modifier [44], which reduces maximum stomatal conductance (and then also GPP) in the Jarvis model and increases RA because of biomass accumulation during forest development.

### 2.3. Virtual stands, model runs and results evaluation

The 3D-CMCC-FEM was first evaluated under observed climate and field data for GPP and NPPwoody (i.e., the NPP for woody compound; gC m–2 year–1) and the diameter at breast height (DBH)(see ‘Model validation’ paragraph in Supplementary Material; [12, 14]). The model was forced with the modeled climate under different emission scenarios, corresponding to the RCP atmospheric CO2 concentration values for the period 1997 to 2100, ranging from 421.4 µmol mol-1 in the ‘best-case scenario’ (RCP2.6) to 926.6 µmol mol-1 of the ‘worst-case scenario’ (RCP 8.5) coming from the ISIMIP-PROFOUND initiative. For comparison purposes, we forced the forest model with a detrended and repeated meteorology and atmospheric CO2 concentration from 1996-2006. The current climate (i.e., no climate change ‘NoCC’) is considered the baseline to compare against climate change scenarios. At the start of the simulations, we created a Composite Forest Matrix (CFM, composed of both measured stand data and “virtual” stand data), following the approach described in Dalmonec et al. [14], to simulate the potential effect of climate stressors on stands of different ages. The 3D-CMCC-FEM has been run at each site to cover the rotation period of each species (from 1997 to 2099) amid the current climate scenario (fixed atmospheric CO2 concentration at the year 2000 of 368.8 μmol mol–1) consisting of detrended and repeated cycles of the present-day observed meteorology from 1996 to 2006 and the Business-as-Usual (BAU) management practices observed at each site (see [28] for the description of BAU applied at each site). Data required to re-initialize the model at every tenth of the rotation length were retrieved from each simulation. Hence, ten additional stands were chosen for each age in the composite matrix and added to the CFM. This collection of virtual forest stands was used to set different starting stand ages at the present day (aget0) due, ideally, to the past silvicultural practice and climate. Under this framework, a landscape of eleven different stands (in age and their relative C-pools and forest structure) for each site is created. These new stands were used, each running from 2006 to 2099, to assess the impact of climate forcing, as the model has already been shown to be sensitive to forest stand development and the relative standing biomass.

The 3D-CMCC-FEM was initialized with the structural attributes of the newly created stands from 1997, which was the starting year of all simulations and for all stands. Modeled climate change simulations under different RCP-emissions scenarios started to differentiate in 2006 (up to 2100). The simulation runs from the different stand initial conditions, corresponding to different aget0 classes, were carried out without forest management as we are interested in the direct climate impact on undisturbed forest stand response, avoiding the confounding effects of forest management on the responses (for forest management effects, see [14]). 825 different simulations were performed as they combined 5 ESMs * 5 RCPs (4 RCPs + 1 current climate scenario) * 11 aget0 classes * 3 sites. Results are reported for MAI (Mean Annual Increment; m3 ha–1 year–1) and TCWS (Total Carbon Woody Stocks; MgC ha-1), respectively, as they are considered some of the most representative and fundamental variables in the carbon cycle and forestry. Following the methodology reported [14] (see Table S1 in Supplementary Materials), we evaluated the model forced with the modeled climate. We compared GPP and NPPwoody against eddy covariance estimates and ancillary data for the years 1997-2005 for DK-Sor and FI-Hyy and 2000-2005 for CZ-BK1. We also compared the diameter at breast height (DBH) in all sites with field measures (see Supplementary Materials).

## 3. Results

Effect of age classes and climate change on total carbon woody stock and increments Norway spruce at CZ-BK1 shows mean TCWS values ranging between ∼70 to ∼140 MgC ha–1 under the NoCC scenario over the century, while from ∼70 to ∼130 MgC ha–1, with a decreasing pattern across all RCPs (Figure 2). In the Norway spruce stands under some ESMs climate forcing (HadGEM2-ES and GFDL-ESM 2M mostly) and under all climate change scenarios, the 3D-CMCC-FEM simulates mortality events for carbon starvation, which increase across stands under gradually warmer climate scenarios and from the oldest stands to the progressively youngest ones.

**Figure 1.**
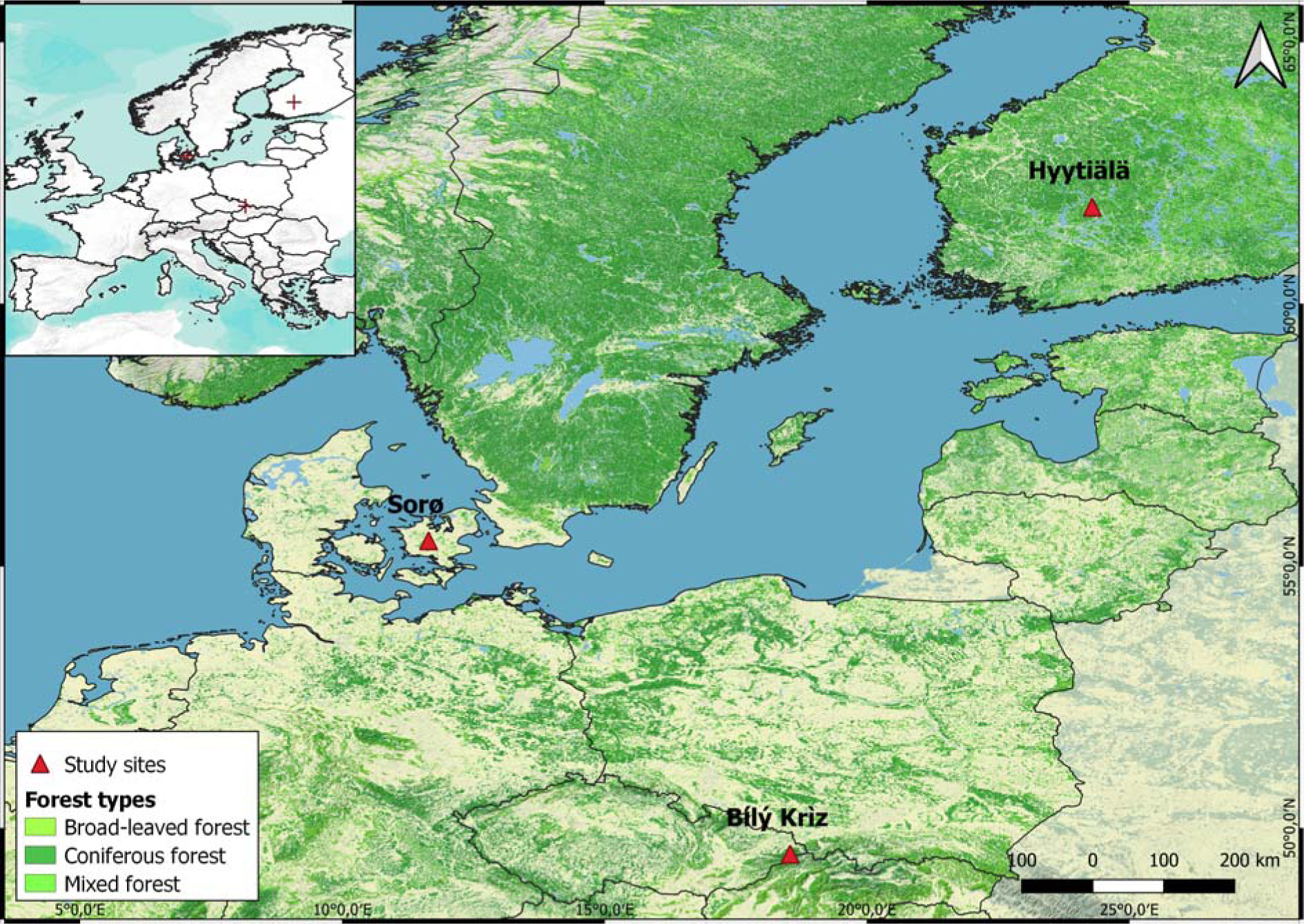
Test site locations in Europe.

**Figure 2.**
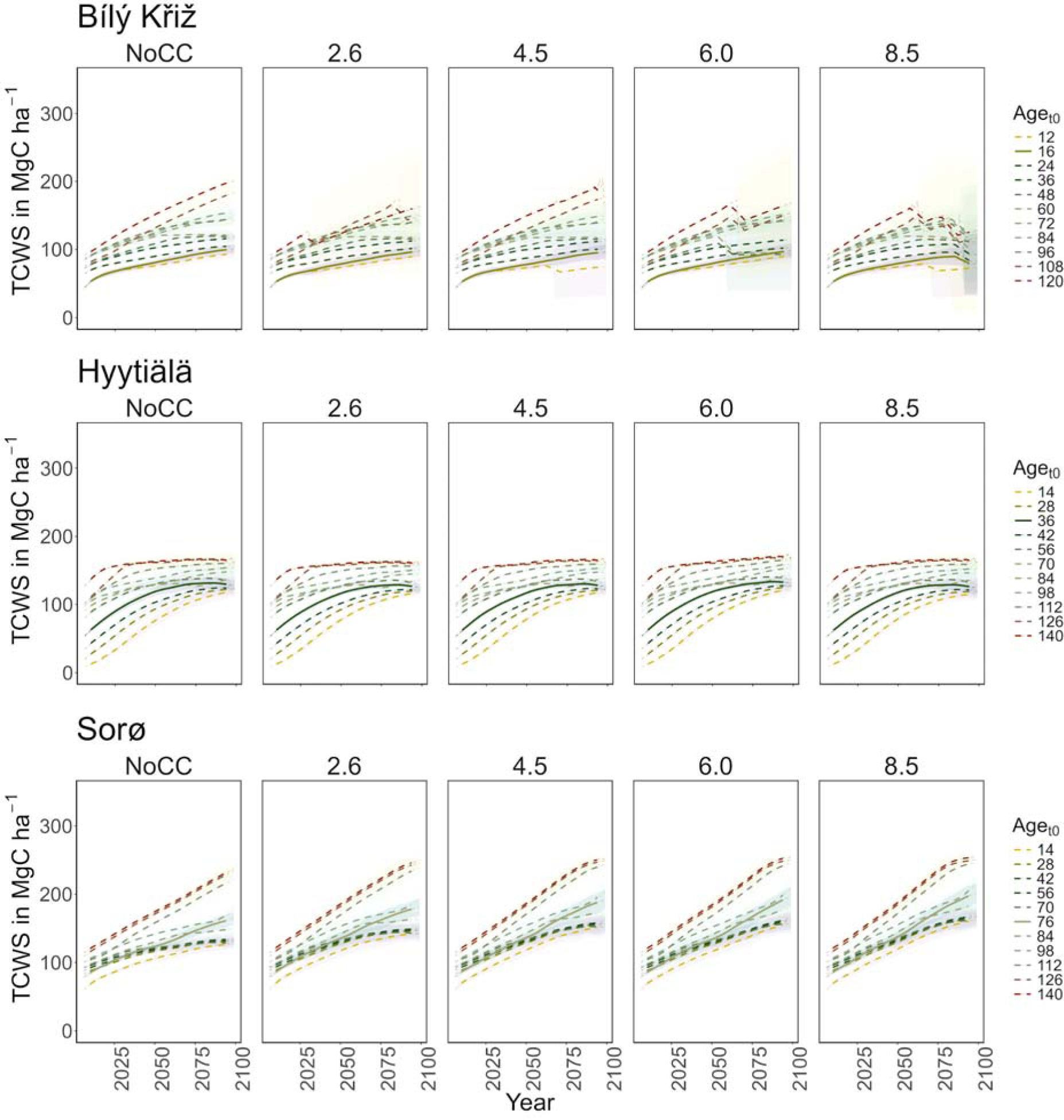
Modeled total carbon woody stock (TCWS) (MgC ha–1) for age classes at the three sites in all scenarios along the simulation period (2006-2099). Lines represent the moving average of 10 years. The solid line corresponds to the real stand, while the dotted lines correspond to the virtual ones. The shaded area represents two standard deviations from the mean predictions with the results from the five ESMs’ climate change scenarios.

Under RCP 8.5, all classes show signs of decay at the end of the century. In the youngest aget0 classes, a sharp decrease in MAI was observed (from 8 to 4 m3 ha–1 year–1), while in the older ones, it holds steady to ∼3 m3 ha–1 year–1 with a peak around 2075 (Figure 3). At FI-Hyy, younger aget0 classes (14-to 42-year-old) showed the fastest increase in TCWS (reaching 120-130 MgC ha– 1 at the end of the century under all scenarios), also reflected in the pattern of MAI. Older aget0 classes showed a more stable trend throughout the simulation (Figure 2), culminating at ∼150 MgC ha–1, with MAI steadily declining from 2.5 to 2 m3 ha–1 year–1. In all scenarios, the Scots pine peaked in the 126 and 56 aget0 in TCWS and MAI, respectively. Minor differences were found in mean TCWS between the NoCC and other RCP scenarios, ranging from –1.6% (140-year-old class under RCP 2.6) to +2.8% (14-year-old class under RCP 6.0). At DK-Sor, results for TCWS show different patterns to other sites, with the highest values ranging between ∼240 MgC ha–1 (under NoCC) to ∼255 MgC ha–1 (under RCP 8.5) at the end of the century with the least TCWS under NoCC. The younger classes showed a shallow increase in TCWS during the simulation period, stabilizing at the end of the century, while the older ones kept growing (Figure 4). DK-Sor was the only site where the tightening of the climate conditions caused a positive effect on the MAI, particularly in the younger classes, reversing the trend from negative to positive at the end of the century.

**Figure 3.**
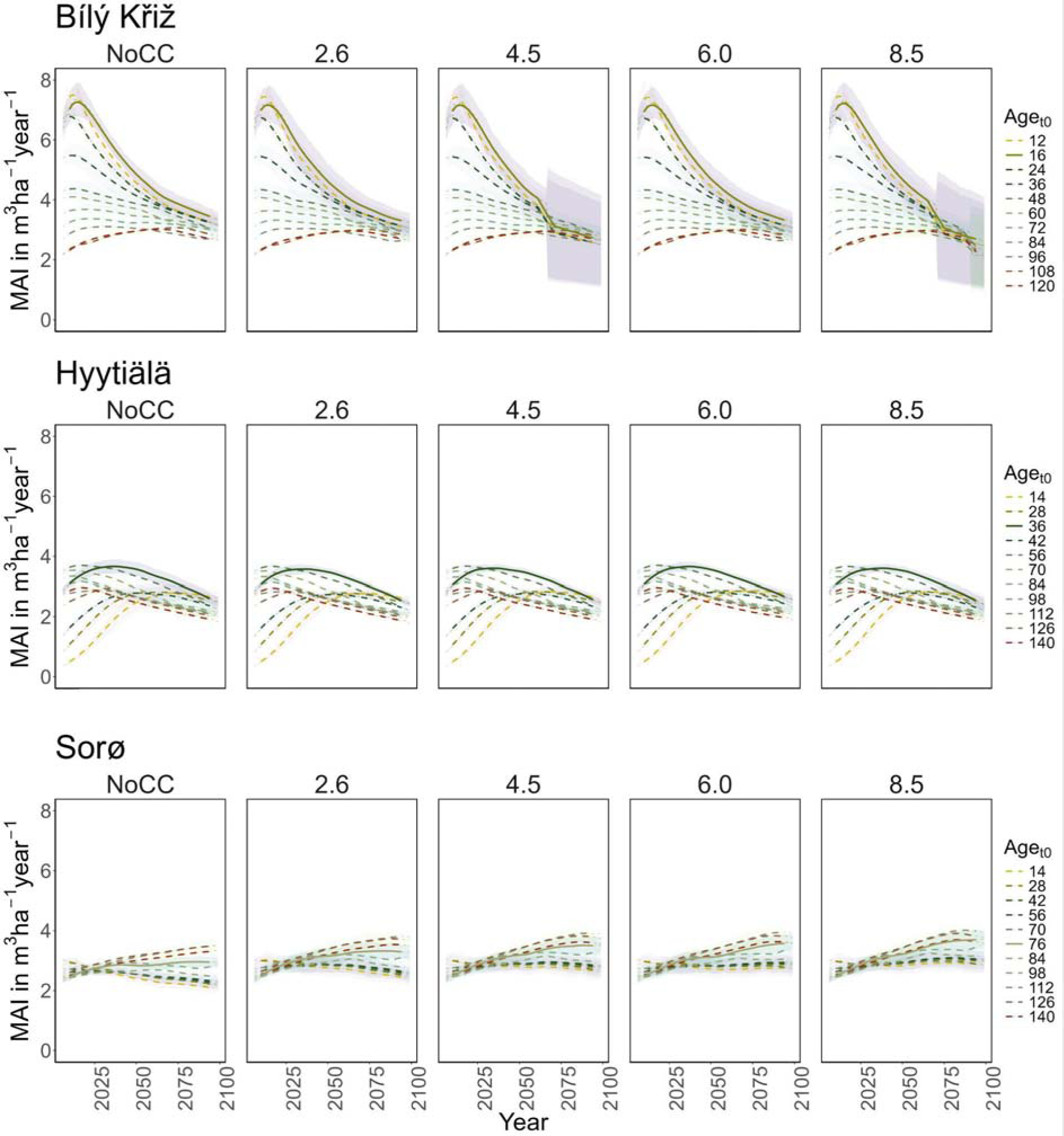
Modeled mean annual increment (MAI) (m3 ha–1 year–1) for age classes at the three sites in all scenarios along the simulation period (2006-2099). Lines represent the moving average of 10 years. The solid line corresponds to the real stand, while the dotted lines correspond to the virtual ones. The shaded area represents two standard deviations from the mean predictions with the results from the five ESMs climate change scenarios.

**Figure 4.**
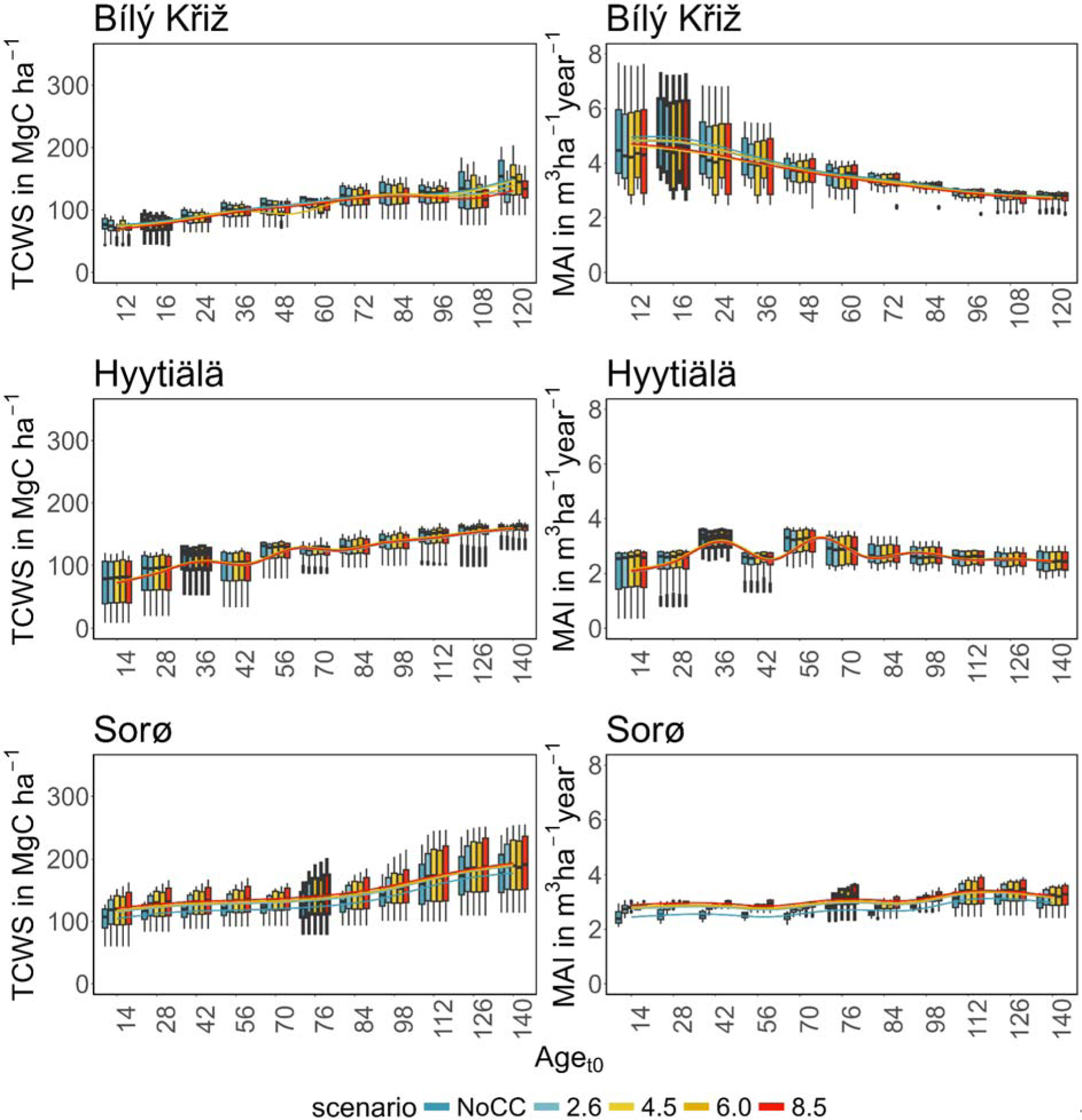
Boxplot of modeled total carbon woody stock (TCWS) (left, MgC ha–1) and mean annual increment (MAI) (right, m3 ha–1 year–1) for age classes at the three sites in the four RCPs scenarios compared to the NoCC (No Climate Change). Boxplots with thick borders correspond to the real stand. Lines are fitted throughout the median of the values of the variables using a generalized additive model.

In summary, a positive growth trend of TCWS over time was found in all sites, with the oldest aget0 classes accounting for the most carbon accumulation. Both conifer stands show a plateau with a reduction in growth at the end of the simulation, which is more pronounced and more severe in the warmest climate scenario. Conversely, the beech stands show a positive growth pattern in all scenarios. Similar results were obtained for MAI, where the conifers showed a decreasing trend over the simulation period despite different magnitudes and patterns among aget0 classes. The beech stands exhibited smaller variations among aget0 than among scenarios concerning other sites. In Table 3, we report the mean value of TCW and MAI over the simulation period for each site and climate scenario.

**Table 1.** Overview of the main site characteristics provided for each forest site. Years of obs. refers to the first and last year of measurement; the temporal resolution of measurement is annual. For the stand values (DBH, Height, BA, Age, Tree density), the range corresponds to the first and last field measurement according to the years of obs. Column. DBH= diameter at breast height; BA= basal area.

| Name | Species | Lat | Long | Aspect | Elevation | Slope | Years of obs. | DBH (cm) | Height (m) | BA (m<sup>2</sup>·h a<sup>-1</sup>) | Age | Tree density (ha<sup>-1</sup>) |
| --- | --- | --- | --- | --- | --- | --- | --- | --- | --- | --- | --- | --- |
| Bílý Kriz | Picea abies | 49.3 | 18.32 | 180 | 875 | 12.5 | 1997–2015 | 8.16–20.47 | 6.26–15.26 | 10.33–36.96 | 16–34 | 2408–1252 |
| Hyytiälä | Pinus sylvestris | 61.85 | 24.29 | 180 | 185 | 2 | 1995–2011 | 15.89–20.58 | 12.61–18.62 | 12.64–18.33 | 34–50 | 870–684 |
| Sorø | Fagus sylvatica | 55.49 | 11.64 | – | 40 | 0 | 1994–2017 | 28.99–48.25 | 24.23–31.15 | 18.50–29.76 | 62–87 | 407–199 |

**Table 2.** Yearly averages of the daily maximum temperature (Tmax), daily minimum temperature (Tmin), daily mean temperature (Tmean), annual precipitation sum (P), daily mean relative humidity (RH), daily mean air pressure (AP), annual sum of global radiation (R, direct + diffuse shortwave radiation) for each of the sites. The column “Years” indicates the data’s acquisition year and the period the average values refer to.

| Site | Source | Years | Tmax (°C) | Tmean (°C) | Tmin (°C) | P (mm) | RH (%) | R (J·cm<sup>-2</sup>) |
| --- | --- | --- | --- | --- | --- | --- | --- | --- |
| Bílý Kriz | Local | 2000–2008 | 11.5 | 7.36 | 3.8 | 1434.56 | 81.99 | 378–774.86 |
| Hyytiälä | Local | 1996–2014 | 7.4 | 4.36 | 1.13 | 604.01 | 77.95 | 309–628.86 |
| Sorø | Local | 1996–2012 | 10.66 | 8.26 | 5.91 | 760.52 | 82.95 | 360–687.83 |

**Table 3.**
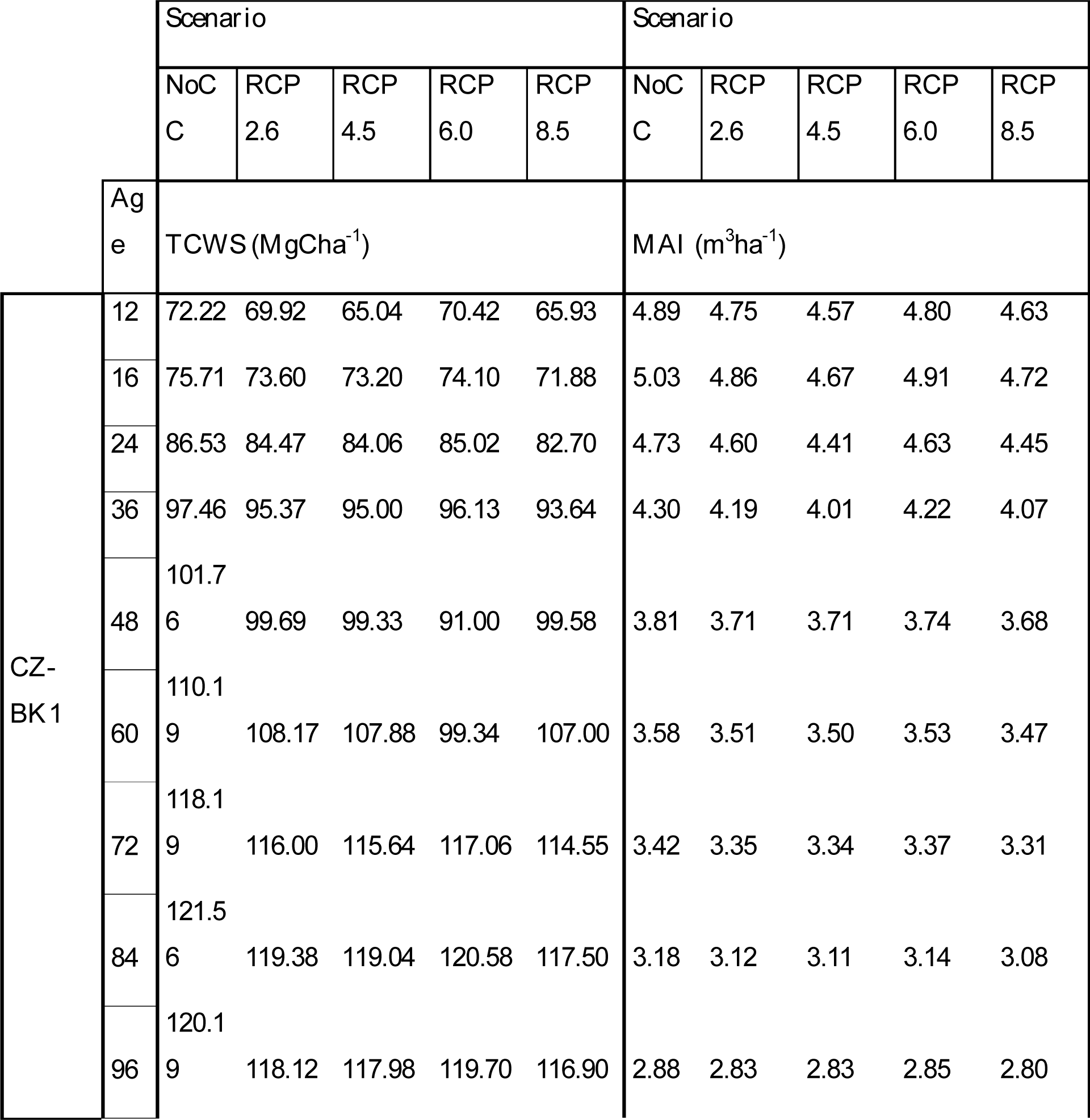
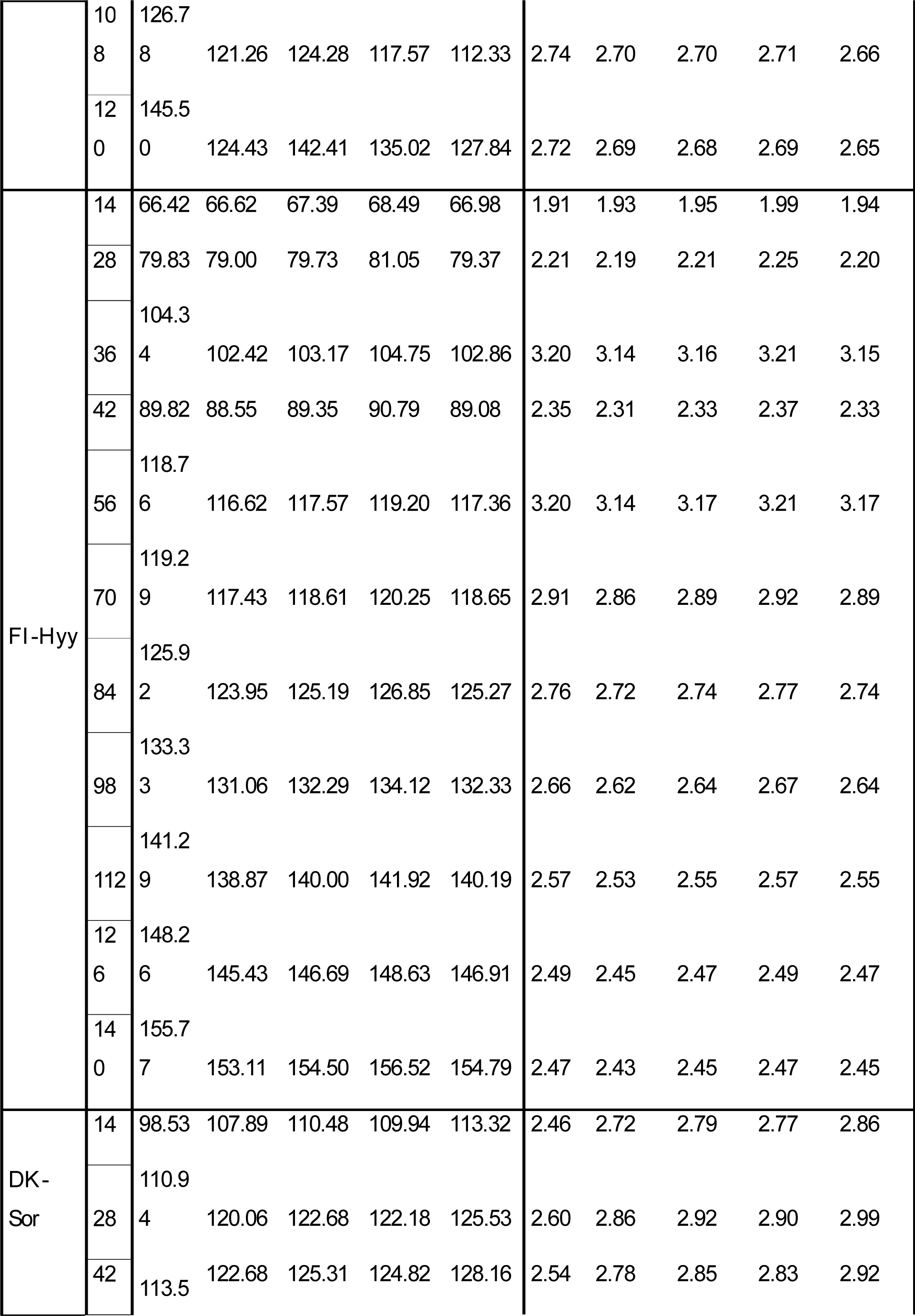
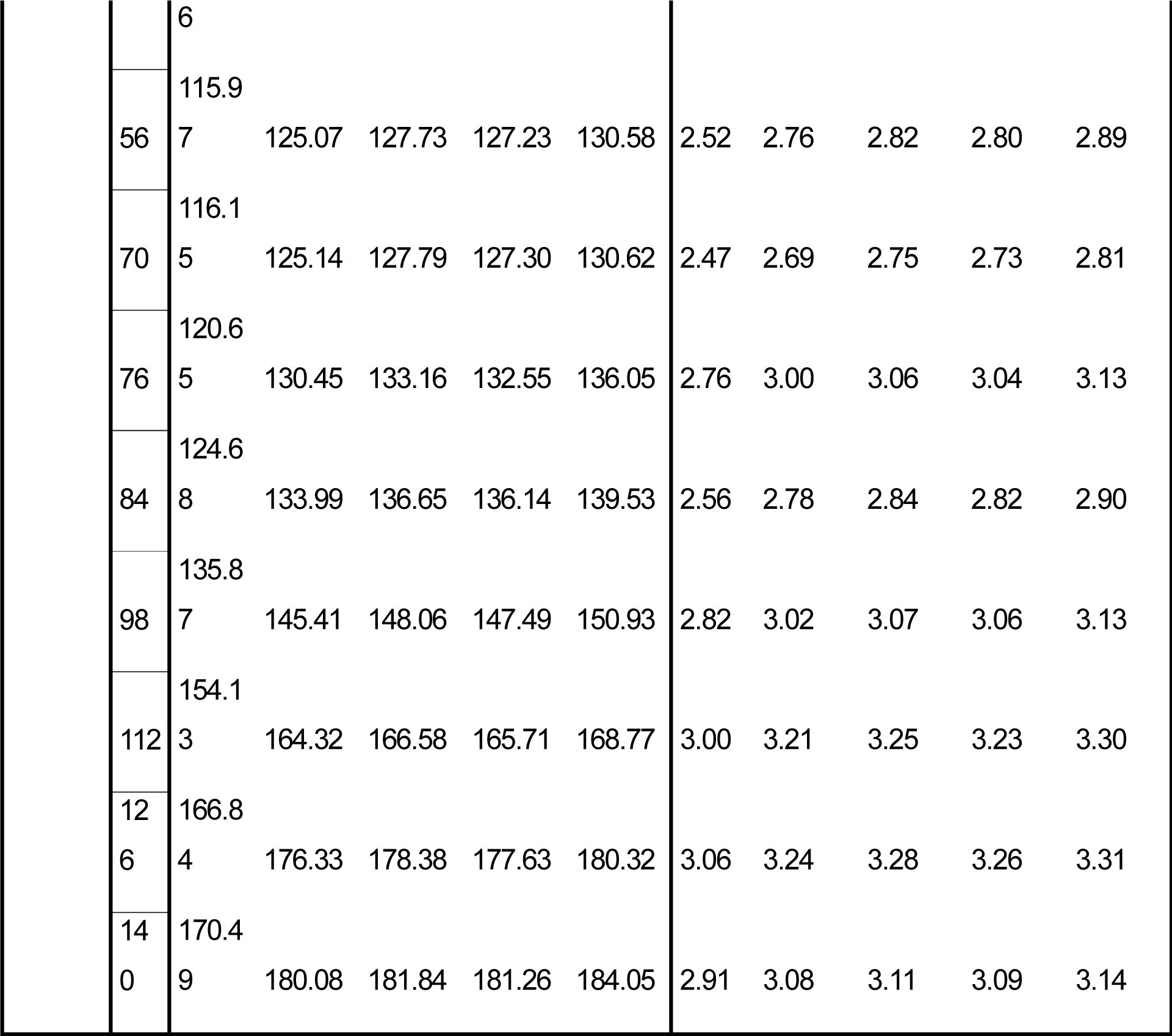
Mean values of total carbon woody stock (TCWS) and mean annual increment (MAI) over the simulation period (2006-2099) for each scenario and age class. CZ-BK1= Bílý Krìz; FI-Hyy= Hyytiälä; DK-Sor= Sorø.

## 4. Discussion

Age-dependent impacts of climate change on forests’ increment and C-stocks The successional stage, represented by forest age, was the main driver controlling C-storage capacity and biomass accumulation, as already known by previous studies [45, 46, 47], with differences greater among different age cohorts under the same scenario than in different climate scenarios under the same age class [12, 14]. The evidence that the carbon budget is mainly controlled by stand age suggests that the effects of climate change on forest cohorts are generally less significant than the effect of age, mainly in terms of the amount of standing biomass. In this sense, age represents multiple and interacting processes, such as tree size [48, 49], forest structural traits (canopy closure and LAI), reduction in stomatal conductance [16], and adaptation to specific environmental conditions which, in turn, make it possible to increases the above-ground biomass (AGB) [50]. The model could reproduce the expected behavior of biomass (and thus carbon) accumulation, simulating rapid growth at a young age and saturation for the oldest age class, but not necessarily at the end of the simulation period. Approaching the physiological optima for the species may benefit the biomass synthesis through an augmented photosynthate supply but may eventually increase the respiratory costs of tissue growth and maintenance despite a strong acclimation capacity [18]. High respiratory costs in warm climates and with low precipitation regimes combined in the older age classes lead to C-starvation and mortality phenomena, as observed for the Norway spruce at the CZ-BK1 site. This indicates that the environment has reached its carrying capacity and that competition for limited resources, such as light and water, is excessively high to sustain more biomass in the oldest age classes.

We found different C-accumulation patterns under climate change between coniferous stands and broadleaves. As expected, increasing temperature and changes in precipitation patterns led to a decline in above-ground biomass in spruce stands, especially in the older age classes. On the contrary, the results show that beech forests at DK-Sor will maintain and even increase C-storage rates under most RCP scenarios. Scots pine forests show an intermediate behavior with a stable stock capacity over time and in different scenarios but with decreasing MAI. These results confirm current observations worldwide that indicate a stronger climate-related decline in conifers forests than in broadleaves [51, 52, 53]. This contrasting response is explained by the different characteristics of the two phyla, in particular, due to the temperature adaptation, with generally lower optimum temperature in conifer and less sensitivity to the length of the growing season.

Similarly, conifers also show lower efficiency in water management because of the shallower root system, which increases the sensitivity to soil aridity and its vulnerability to drought events [54]. Recent studies confirm that growth decline is more pronounced in conifers than broadleaf, especially beech forests, in the most northern species distribution [55]. Our results confirm the same growth patterns found by recent studies [47, 53, 56], where broadleaves outperform conifers in productivity, and climate warming will probably exacerbate these opposite growth patterns.

However, despite some studies suggesting that age modulates different adaptation strategies to some extent, it remains unclear whether younger trees may be more affected by climate change than older ones. Bennett et al. [57], in a global analysis, found that droughts consistently had more severe impacts on larger (older) trees, while Wang et al. [11] observed a more substantial and sharper decline in basal area increment in young Korean pine in China. Hogg et al. [58] found that the percentage decrease in biomass growth was not significantly different for young, productive stands compared to older, less productive ones. Our study suggests that warmer and drier conditions and extended growing seasons will affect younger stands more than older ones, but with different trends among species. In particular, MAI will be positively affected in younger beech forests, while it will remain stable in older stands. On the contrary, climate change will strongly impact the growth rate of young conifers stands more than older ones. Older forests tend to be more stable and resilient than younger ones due to their rugged and stable interaction with climate triggers and better responsiveness to environmental changes. The year-to-year climate variability is buffered by larger carbon pools in sapwood and reservoirs in older trees, leading to higher long-term stability than younger trees [12]. In this sense, ages represent the “memory” of the forest to past climate and disturbance regimes, which align the species-specific traits to the environmental conditions in which they grow, creating the niches in which AGB accumulates [52, 59].

Despite numerous efforts to decipher forests’ response to climate change, the intricate methods employed by tree species to withstand extreme climates still need to be fully unveiled. Further research exploiting ecophysiological models explicitly accounting for age, tree-ring experiments, and remote sensing will be critical to understanding forest ecosystems’ adaptation strategies to climate change, particularly in the face of rapid warming and extreme disturbances. A better understanding of the interaction between forests and climate can inform better forest management strategies, ultimately dampening the impacts of climate change on forest ecosystems.

## 5. Limitations

The presented modeling framework has some limitations that should be considered. Firstly, natural disturbances as consequences of climate change, such as windstorms, forest fires, and insect outbreaks, were not simulated. These disturbances cause changes in carbon stocks, nutrients, and soil conditions and contribute to the global release of CO2 in the atmosphere, ultimately leading to increasing temperature and radiation. In contrast, climate extreme events are considered to be already included in the climate scenarios used to force the model and, thus, already accounted for in the model outputs. Additionally, other indirect alterations due to climate change of key drivers, such as nitrogen deposition, phosphorus, or ozone, which can somewhat amplify or reduce our results, were not assessed. Nonetheless, some studies (e.g., [60]) lend credence to the notion that this phenomenon may not be applicable across the board. They highlight the significant responsiveness of various tree species to CO2 fertilization across a wide range of nutrient availability. Finally, no allowance was made for the possibility of species migration to and from the study areas. However, these dynamics may require longer timescales than those simulated in this study.

## 5. Conclusions

Forest age is confirmed to be a significant factor in determining the carbon storage capacity and biomass accumulation in forest ecosystems, especially in the context of future climate uncertainty. The effects of species, site location, stand-level characteristics, and development stage vary significantly and are contingent on specific factors. We observed that differences in biomass accumulation were more pronounced among different age cohorts than among different climate scenarios within the same age class, with contrasting carbon accumulation patterns under climate change between coniferous and broadleaf forests. Furthermore, our findings shed light on the differential impacts of climate change on younger versus older forest stands. Warmer and drier conditions are projected to affect younger stands more severely, particularly in coniferous forests. However, older forests will likely exhibit greater stability and resilience due to their accumulated carbon pools and enhanced adaptability to environmental changes. While our study provides valuable insights, it also underscores the need for further research to unravel the complex mechanisms by which forests adapt to climate change. This deeper understanding can inform more effective forest management strategies, helping to mitigate the impacts of climate change on forest ecosystems in the future. The varying responses of different tree species highlight the need for tailored management approaches and conservation efforts to enhance the resilience of our forests.

## Author Contributions

E.V.: Data curation, Formal analysis, Investigation, Writing – original draft, Writing – review & editing; D.D.: Data curation, Formal analysis, Investigation, Writing – review & editing; M.M.: Writing – review & editing; E.G.: Writing – review & editing; F.G.: Writing – review & editing; G.D.: Writing – review & editing; M.N.: Writing – review & editing; G.C.: Writing – review & editing; A.C.: Formal analysis, Investigation, Writing – original draft, Writing – review & editing, Conceptualization. All authors have read and agreed to the published version of the manuscript.

## Funding

“FORESTNAVIGATOR” Horizon Europe research and innovation program under grant agreement No. 101056875; Research Projects of National Relevance funded by the Italian Ministry of University and Research entitled: “Multi-scale observations to predict Forest response to pollution and climate change” (MULTIFOR, project number 2020E52THS) Data Availability Statement: The 3D-CMCC-FEM v.5.6 model code is publicly available and can be found on the GitHub plat-form at: https://github.com/Forest-Modelling-Lab/3D-CMCC-FEM). The raw data supporting the conclusions of this article will be made available by the authors on request.

## Conflicts of Interest

The authors declare no conflicts of interest.

## Acknowledgments

E.V. and A.C. acknowledge funding by the project “FORESTNAVIGATOR” Horizon Europe research and innovation program under grant agreement No. 101056875. M.M., E.G., F.G., and A.C. acknowledge funding by the project “OptForEU” Horizon Europe research and innovation program under grant agreement No. 101060554. D.D. and A.C. also acknowledge the project funded under the National Recovery and Resilience Plan (NRRP), Mission 4 Component 2 Investment 1.4 - Call for tender No. 3138 of 16 December 2021, rectified by Decree n.3175 of 18 December 2021 of Italian Ministry of University and Research funded by the European Union – NextGenerationEU under award Number: Project code CN_00000033, Concession Decree No. 1034 of 17 June 2022 adopted by the Italian Ministry of University and Research, CUP B83C22002930006, Project title “National Biodiversity Future Centre - NBFC”. E.V. and A.C. also acknowledge funding from the MIUR Project (PRIN 2020) “Unraveling interactions between WATER and carbon cycles during drought and their impact on water resources and forest and grassland ecosySTEMs in the Mediterranean climate (WATERSTEM)” (Project num-ber: 20202WF53Z), “WAFER” at CNR (Consiglio Nazionale delle Ricerche), and by PRIN 2020 (cod 2020E52THS) - Research Projects of National Relevance funded by the Italian Ministry of University and Research entitled: “Multi-scale observations to predict Forest response to pollution and climate change” (MULTIFOR, project number 2020E52THS). We also thank the ISIMIP project (https://www.isimip.org/) and the COST Action FP1304 “PROFOUND” (Towards Robust Projections of European Forests under Climate Change), supported by COST (European Cooperation in Science and Technology) for providing us the climate historical scenarios and site data used in this work. This work used eddy covariance data acquired and shared by the “FLUXNET” community, including these networks: AmeriFlux, AfriFlux, AsiaFlux, CarboAfrica, CarboEurope-IP, CarboItaly, CarboMont, ChinaFlux, Fluxnet-Canada, GreenGrass, ICOS, KoFlux, LBA, NECC, OzFlux-TERN, TCOS-Siberia, and USCCC. The ERA-Interim reanalysis data are pro-vided by ECMWF and processed by LSCE. The FLUXNET eddy covari-ance data processing and harmonization was carried out by the European Fluxes Database Cluster, AmeriFlux Management Project, and Fluxdata Project of FLUXNET, with the support of CDIAC and ICOS Ecosystem Thematic Center, and the OzFlux, ChinaFlux, and AsiaFlux offices. We acknowledge the World Climate Research Programme’s Working Group on Coupled Modelling, which is responsible for CMIP, and we thank the respective climate modeling groups for producing and making available their model output. The U.S. Department of Energy’s Program for Climate Model Diagnosis and Intercomparison at Lawrence Livermore National Laboratory provides coordinating support for CMIP and led the development of software infrastructure in partnership with the Global Organization for Earth System Science Portals.

## Supplementary materials Model evaluation

Daily model outputs were evaluated against eddy-covariance and measured struc-tural data at the site level, in terms of percentage root mean squared error (RMSE%) and Pears’s correlation coefficient (r). The GPP evaluation for simulations forced with observed site-specific daily weather data (1997-2005 for FI-Hyy, and DK-Sor, and 2000-2005 for CZ-BK1) resulted in an RMSE% of 1.05, 1.52, and 1.43, with r values of 0.92, 0.87 and 0.94 for FI-Hyy, CZ-BK1 and DK-Sor, respectively (Table 1). Similar results were obtained for NPPwoody in the site of DK-Sor and CZ-BK1 (351 ±61 gC m–2 year–1 vs. 346 ±36 gC m–2 year–1 measured, and 442 ±79 gC m–2 year–1 vs. 380 ± 38 gC m–2 year–1 measured, respectively). At FI-Hyy, modeled NPPwoody data was overestimated in respect to the measured values (317 ±21 gC m–2 year–1 vs. 228 ±23 gC m–2 year–1 measured).

**Table S1:**
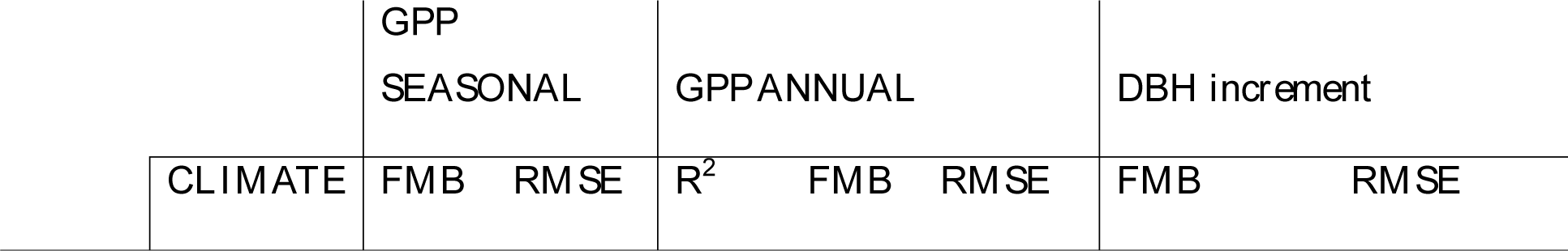
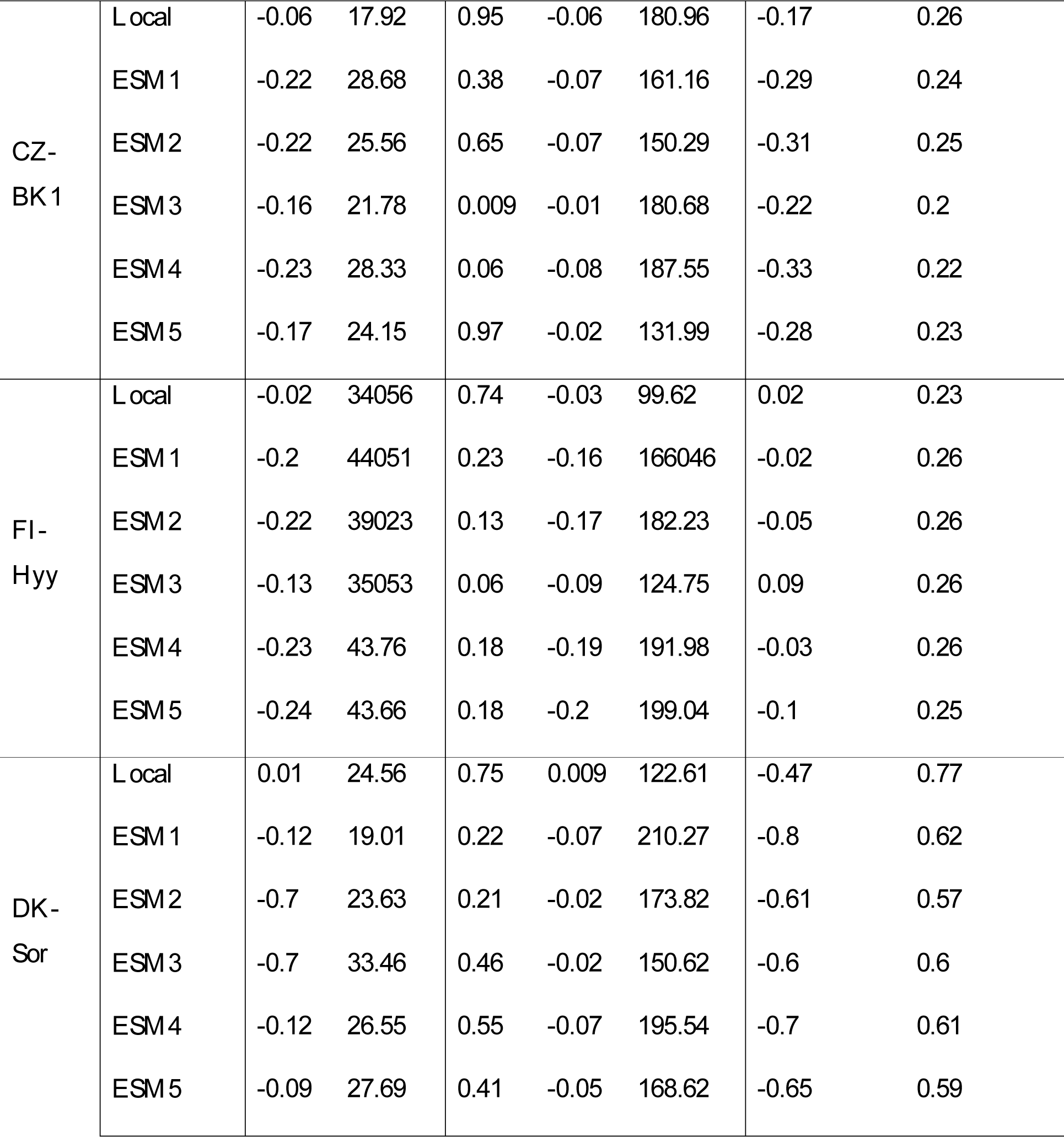
Performance statistics (coefficient of determination R2, relative root mean square error RMSE (gC m–2 day–1) and Fractional Mean Bias, FMB) computed from monthly seasonal values and annual series of model gross primary productivity, GPP, against eddy covariance estimated and diametric annual increment data, DBH increment, against measured data. Results are reported for simulations forced with local and modeled climate (i.e., ESM) (ESM1, 2, 3, 4, 5 refer to HadGEM2-ES, IPSL-CM5A-LR, MIROC-ESM-CHEM, GFDL-ESM 2M, and NorESM1-M, respectively).

|  | GPP SEASONAL |  | GPP ANNUAL |  |  | DBH increment |  |
| --- | --- | --- | --- | --- | --- | --- | --- |
| CLIMATE | FMB | RMSE | R<sup>2</sup> | FMB | RMSE | FMB | RMSE |

|  |  |  |  |  |  |  |  |  |
| --- | --- | --- | --- | --- | --- | --- | --- | --- |
| **CZ-BK1** | **Local** | -0.06 | 17.92 | 0.95 | -0.06 | 180.96 | -0.17 | 0.26 |
|  | **ESM1** | -0.22 | 28.68 | 0.38 | -0.07 | 161.16 | -0.29 | 0.24 |
|  | **ESM2** | -0.22 | 25.56 | 0.65 | -0.07 | 150.29 | -0.31 | 0.25 |
|  | **ESM3** | -0.16 | 21.78 | 0.009 | -0.01 | 180.68 | -0.22 | 0.2 |
|  | **ESM4** | -0.23 | 28.33 | 0.06 | -0.08 | 187.55 | -0.33 | 0.22 |
|  | **ESM5** | -0.17 | 24.15 | 0.97 | -0.02 | 131.99 | -0.28 | 0.23 |
| **FI-Hyy** | **Local** | -0.02 | 34056 | 0.74 | -0.03 | 99.62 | 0.02 | 0.23 |
|  | **ESM1** | -0.2 | 44051 | 0.23 | -0.16 | 166046 | -0.02 | 0.26 |
|  | **ESM2** | -0.22 | 39023 | 0.13 | -0.17 | 182.23 | -0.05 | 0.26 |
|  | **ESM3** | -0.13 | 35053 | 0.06 | -0.09 | 124.75 | 0.09 | 0.26 |
|  | **ESM4** | -0.23 | 43.76 | 0.18 | -0.19 | 191.98 | -0.03 | 0.26 |
|  | **ESM5** | -0.24 | 43.66 | 0.18 | -0.2 | 199.04 | -0.1 | 0.25 |
| **DK-Sor** | **Local** | 0.01 | 24.56 | 0.75 | 0.009 | 122.61 | -0.47 | 0.77 |
|  | **ESM1** | -0.12 | 19.01 | 0.22 | -0.07 | 210.27 | -0.8 | 0.62 |
|  | **ESM2** | -0.7 | 23.63 | 0.21 | -0.02 | 173.82 | -0.61 | 0.57 |
|  | **ESM3** | -0.7 | 33.46 | 0.46 | -0.02 | 150.62 | -0.6 | 0.6 |
|  | **ESM4** | -0.12 | 26.55 | 0.55 | -0.07 | 195.54 | -0.7 | 0.61 |
|  | **ESM5** | -0.09 | 27.69 | 0.41 | -0.05 | 168.62 | -0.65 | 0.59 |

**Figure S1:**
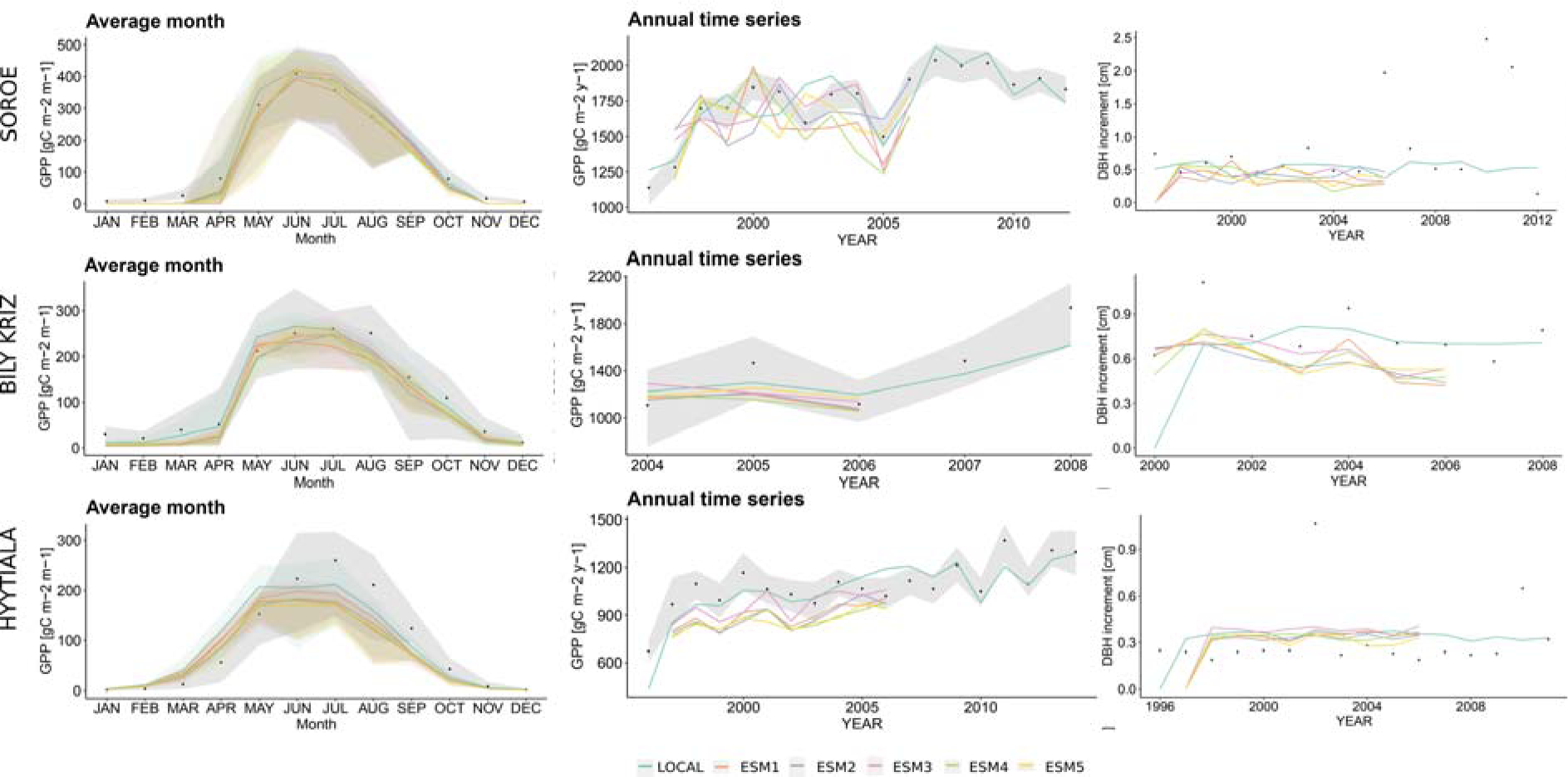
Evaluation of monthly seasonal GPP (gC m–2 month–1) fluxes (left column) and annual (gC m–2 year–1) fluxes (central column) for the sites of Sorø, Bily Kriz, and Hyytiala (rows). Quali-ty-checked and -filtered GPP values evaluated at the sites by the eddy covariance technique are reported as black dots. The shaded area for seasonal values reports the maximum and minimum monthly values recorded in the time series. The shaded area for annual data represents the relative uncertainty bounds. In the third column, a comparison of the predicted annual DBH increment (cm y-1) with site observations at the three sites is reported. Measured data are shown as black dots. Simulated data are reported as continuous lines.

## Notes

### Competing Interest Statement

The authors have declared no competing interest.

### Summary of Updates

Added two tables in material and methods section; Updated figure 1. Minor spelling and typos correction

## References

1. Favero, A.; Mendelsohn, R.; Sohngen, B.; Stocker, B. Assessing the long-term interactions of climate change and timber markets on forest land and carbon storage. Environ. Res. Lett. 2021, 16 014051 10.1088/1748-9326/abd589

2. Vangi, E.; D’Amico, G.; Francini, S.; Giannetti, F.; Lasserre, B.; Marchetti, M.; McRoberts, R.E.; Chirici, G. The effect of forest mask quality in the wall-to-wall estimation of growing stock volume. Rem. Sens. 2021, 13, 1038. 10.3390/rs13051038

3. Noce, S.; Collalti, A.; Valentini, R.; & Santini, M. Hot spot maps of forest presence in the Mediterranean basin. iFor-est-Biogeosciences and Forestry 2016, 9(5), 766.

4. Lionello, P.; Scarascia, L. The relation between climate change in the Mediterranean region and global warming. Reg. Environ. Chang. 2018, 18, 1481–1493. 10.1007/s10113-018-1290-1

5. Pretzsch, H.; Biber, P.; Schütze, G. et al. Forest stand growth dynamics in Central Europe have accelerated since 1870. Nat Commun 2014, 5, 4967. 10.1038/ncomms5967

6. He Y.; Liu Y.; Lei L.; Terrer C.; Huntingford C.; Peñuelas J.; Xu H.; Piao, S. CO2 fertilization contributed more than half of the observed forest biomass increase in northern extra-tropical land. Global Change Biology, 2023, 00, 1–14. 10.1111/gcb.16806

7. Roebroek, Caspar T.J.; et al. “Releasing global forests from human management: How much more carbon could be stored?.” Science 2023, 380.6646: 749–753.

8. Duffy, K.A.; Schwalm, C.R.; Arcus, V.L.; Koch, G.W.; Liang, L.L.; Schipper, L.A. How close are we to the temperature tipping point of the terrestrial biosphere? Sci Adv. 2021 Jan 13;7(3):eaay1052.

9. Nabuurs, G.J.; Verkerk, P.J.; Schelhaas, M.J.; González, O.J.R.; Trasobares, A.; Cienciala, E. Climate-Smart Forestry: mitigation impacts in three European regions. From Science to Policy 2018, 6. European Forest Institute.

10. Wang, X.; Pederson, N.; Chen, Z.; Lawton, K.; Zhu, C.; Han, S. Recent rising temperatures drive younger and southern Korean pine growth decline. Science of the total environment 2019, 649, 1105–1116.

11. Gregor, K.; Krause, A.; Reyer, C.P.O. et al. Quantifying the impact of key factors on the carbon mitigation potential of man-aged temperate forests. Carbon Balance Manage 2024, 19, 10. 10.1186/s13021-023-00247-9

12. Vangi, E.; Daniela, D.; Elisa, C.; Gina, M.; Leonardo, B.; Paulina, F.P.; Elisa, G.; Alessandro, C.; Andrea, C.; Gherardo, C.; Alessio, C. Stand age diversity and climate change affect forests’ resilience and stability, although unevenly. bioRxiv 2023.07.12.548709; doi: 10.1101/2023.07.12.548709

13. Erb, K-H.; Haberl, H.; Le Noë, J.; Tappeiner, U.; Tasser, E.; Gingrich, S. Changes in perspective needed to forge ‘no-regret’ for-est-based climate change mitigation strategies. GCB Bioenergy. 2022, 14(3):246–57. 10.1111/gcbb.12921.

14. Dalmonech, D.; Marano, G.; Amthor, J.; Cescatti, A.; Lindner, M.; Trotta, C.; Collalti, A. Feasibility of enhancing carbon se-questration and stock capacity in temperate and boreal European forests via changes to forest management. Agricultural and Forest Meteorology 2022, 327(109203), 10.1016/j.agrformet.2022.109203

15. Anderson-Teixeira, K.J.; Miller, A.D.; Mohan, J.E.; Hudiburg, T.W.; Duval, B.D.; Delucia, E.H. Altered dynamics of forest recovery under a changing climate. Global Change Biology, 2013, 19, 10.1111/gcb.12194

16. Ryan, M.G., Binkley, D.; Fownes, J.H. Age-related decline in forest productivity: pattern and process. Advances in ecological re-search 1997, 27, 213–262. 10.1016/S0065-2504(08)60009-4

17. Goulden, M.L.; McMillan, A.M.; Winston, G.C.; Rocha, A.V.; Manies, K.L.; Harden, J.W.; Bond-Lamberty, B.P. Patterns of NPP, GPP, respiration, and NEP during boreal forest succession. Global Change Biology 2011, 17(2), 855–871. 10.1111/j.1365-2486.2010.02274.x

18. Collalti A.; Tjoelker M.G.; Hoch G.; Mäkelä A.; Guidolotti G.; Heskel M.; Petit G.; Ryan M.G.; Battipaglia G.; Prentice I.C. Plant respiration: Controlled by photosynthesis or biomass?. Global Change Biology 2020, 26(3): 1739–1753 10.1111/gcb.14857

19. Collalti, A.; Ibrom, A.; Stockmarr, A.; Cescatti, A.; Alkama, R.; Fernandez-Martínez, M.; Matteucci, G.; Sitch, S.; Friedlingstein, P.; Ciais, P.; Goll, D. S.; Nabel, J. E. M. S.; Pongratz, J.; Arneth, A.; Haverd, V.; Prentice, I. C. Forest production efficiency increases with growth temperature. Nature Communications. Nature Communications, 2020, 11: 5322, 10.1038/s41467-020-19187-w

20. FOREST EUROPE, 2018: State of Europe’s Forests 2018.

21. FOREST EUROPE, 2020: State of Europe’s Forests 2020.

22. Collalti, A.; Marconi, S.; Ibrom, A.; Trotta, C.; Anav, A.; D’Andrea, E.; Matteucci, G.; Montagnani, L.; Gielen, B.; Mammarella, I.; Grünwald, T.; Knohl, A.; Berninger, F.; Zhao, Y.; Valentini, R.; Santini, M. Validation of 3D-CMCC Forest Ecosystem Model (v.5.1) against eddy covariance data for ten European forest sites. Geoscientific Model Development 2016, 9, 1-26, 10.5194/gmd-9-479-2016

23. Collalti, A.;, et al. Monitoring and Predicting Forest Growth and Dynamics, CNR Edizioni, 2024, DOI: 10.32018/ForModLab-book-2024

24. Collalti, A.; Trotta, C.; Keenan, T.F.; Ibrom, A.; Bond-Lamberty, B.; Grote, R.; Vicca, S.; Reyer, C.P.O.; Migliavacca, M.; Ver-oustraete, F.; Anav, A.; Campioli, M.; Scoccimarro, E.; ̌Sigut, L.; Grieco, E.; Cescatti, A.; Matteucci, G. Thinning can reduce losses in carbon use efficiency and carbon stocks in managed forests under warmer climate. J. Adv. Model. Earth Syst. 2018, 10 (10), 2427–2452. 10.1029/2018MS001275

25. Mahnken, M.; Cailleret, M.; Collalti, A.; Trotta, C., Biondo, C.; D’Andrea, E.; Dalmonech, D.; Marano, G.; Mäkelä, A.; Minunno, F.; Peltoniemi, M.; Trotsiuk, V.; Nadal-Sala, D.; Sabate, S.; Vallet, P.; Aussenac, R.; Cameron, D.R.; Bohn, F.J.; Grote, R.; Reyer, C.P.O. Accuracy, realism and general applicability of European forest models. Glob. Change Biol. 2022, 00, 1–23. 10.1111/gcb.16384.

26. Moss, R.; Edmonds, J.; Hibbard, K. The next generation of scenarios for climate change research and assessment. Nature 2010, 463, 747–756

27. van Vuuren, D.; Edmonds, J.; Kainuma, M.; Riahi, K.; Thomson, A.; Hibbard, K.;, et al. The representative concentration path-ways: An overview. Climatic Change 2011, 109(1-2), 5–31. 10.1007/s10584-011-0148-z

28. Reyer, C.P.O.; Silveyra Gonzalez, R.; Dolos, K.; Hartig, F.; Hauf, Y.; Noack, M.; Lasch-Born, P.; Rötzer, T.; Pretzsch, H.; Meesenburg, H.; Fleck, S.; Wagner, M.; Bolte, A.; Sanders, T.G.M.; Kolari, P.; Mäkelä, A.; Vesala, T.; Mammarella, I.; Pumpanen, J.; Frieler, K. The PROFOUND Database for evaluating vegetation models and simulating climate impacts on European forests. Earth Syst. Sci. Data 2020, 12 (2), 1295–1320. 10.5194/essd-12-1295-2020

29. Vacchiano, G.; Magnani, F.; Collalti A. Modeling Italian forests: state of the art and future challenges. iForest 2012, 5: 113–120. - doi: 10.3832/ifor0614-005

30. Collalti, A.; Perugini, L.; Santini, M.; Chiti, T.; Nolè A.; Matteucci, G.; Valentini, R. A process-based model to simulate growth in forests with complex structure: Evaluation and use of 3D-CMCC Forest Ecosystem Model in a deciduous forest in Central Italy. Ecological Modelling 2014, 272, 362– 378, 10.1016/j.ecolmodel.2013.09.016

31. Farquhar, G.; von Caemmerer, S.; and Berry, J. A biogeochemical model of photosynthetic CO2 assimilation in leaves of C3 species. Planta, 1980, 149, 78–90.

32. de Pury, D. G. G.; Farquhar, G. D. Simple scaling of photosynthesis from leaves to canopies without the errors of bigCleaf models. Plant, Cell and Environment 1997, 20(5), 537–557. 10.1111/j.1365-3040.1997.00094.x

33. Bernacchi, C. J.; Singsaas, E. L.; Pimentel, C.; Portis Jr, A. R.; Long, S. P. Improved temperature response functions for models of RubiscoClimited photosynthesis. Plant, Cell and Environment 2001, 24(2), 253–259. 10.1111/j.1365-3040.2001.00668.x

34. Bernacchi, C. J.; Calfapietra, C.; Davey, P. A.; Wittig, V. E.; ScarasciaCMugnozza, G. E.; Raines, C. A.; Long, S. P. Photosynthesis and stomatal conductance responses of poplars to freeCair CO2 enrichment (PopFACE) during the first growth cycle and im-mediately following coppice. New Phytologist 2003, 159(3), 609–621. 10.1046/j.1469-8137.2003.00850.x

35. Kattge, J.; Knorr, W. Temperature acclimation in a biochemical model of photosynthesis: a reanalysis of data from 36 spe-cies. Plant, cell & environment 2007, 30(9), 1176–1190.

36. McDowell, N. Mechanism linking drought, hydraulics, carbon metabolism, and vegetation mortality. Plant Physiology 2011, 155, 1051–1059.

37. Rowland, L.; da Costa, A. C. L.; Galbraith, D. R.; Oliveira, R. S.; Binks, O. J.; Oliveira, A. A. R.;, et al. Death from drought in tropical forests is triggered by hydraulics not carbon starvation. Nature 2015, 528, 119–122

38. Collalti, A.; Biondo, C.; Buttafuoco, G.; Maesano, M.; Caloiero, T.; Lucà, F.; Pellicone, G.; Ricca, N.; Salvati, R.; Veltri, A.; Scara-scia-Mugnozza, G.; Matteucci, G. Simulation, calibration and validation protocols for the model 3D-CMCC-CNR-FEM: a case study in the Bonis’ watershed (Calabria), Italy. Forest@ 2017, 14(14): 247–256, 10.3832/efor2368-014

39. Merganičová, K.; Merganič, J.; Lehtonen, A.; Vacchiano, G.; Sever, M. Z. O.; Augustynczik A.L.D.; Grote R.; Kyselová I.; Mäkelä A.; Yousefpour R.; Krejza J.; Collalti A.; Reyer C.P.O. Forest carbon allocation modelling under climate change. Tree Physiology 2019, 39(12), 1937–1960. 10.1093/treephys/tpz105

40. Kira, T.; Shidei, T. Primary production and turnover of organic matter in different forest ecosystems of the western Pacific. Japanese Journal of Ecology 1967, 17(2), 70–87.

41. Odum, E. P. The Strategy of Ecosystem Development: An understanding of ecological succession provides a basis for resolv-ing man’s conflict with nature. Science 1969, 164(3877), 262-270.

42. Tang, J.; Luyssaert, S.; Richardson, A. D.; Kutsch, W.; Janssens, I. A. Steeper declines in forest photosynthesis than respiration explain age-driven decreases in forest growth. Proceedings of the National Academy of Sciences 2014, 111(24), 8856–8860. 10.1073/pnas.1320761111

43. Mäkelä, A.; Valentine, H. The ratio of NPP to GPP: evidence of change over the course of stand development. Tree Physiology 2000, 21:1015–1030. 10.1093/treephys/21.14.1015

44. Landsberg, J. J.; Waring, R. H. A generalised model of forest productivity using simplified concepts of radiation-use efficiency, carbon balance and partitioning. Forest ecology and management 1997, 95(3), 209–228. 10.1016/S0378-1127(97)00026-1

45. Anderson, K. J.; Allen, A. P.; Gillooly, J. F.; Brown, J. H. Temperature-dependence of biomass accumulation rates during sec-ondary succession. Ecol. Lett. 2006, 9 673–82

46. Cook-Patton, S.C.;, et al. Mapping carbon accumulation potential from global natural forest regrowth. Nature 2020, 585 545–50

47. Anderson-Teixeira, K.J.; Herrmann, V.; Banbury Morgan, R. et al; Carbon cycling in mature and regrowth forests globally. Environmental Research Letters 2021, 16 (5). 053009. ISSN 1748-9326 10.1088/1748-9326/abed01

48. Ouyang, S.; Xiang, W.; Wang, X.; Xiao, W.; Chen, L.; Li, S.;, et al. Effects of stand age, richness and density on productivity in subtropical forests in China. J. Ecol. 2019, 107, 2266–2277. doi: 10.1111/1365-2745.13194

49. Ullah, F.; Gilani, H.; Sanaei, A.; Hussain, K.; Ali, A. Stand structure determines aboveground biomass across temperate forest types and species mixture along a local-scale elevational gradient. Forest Ecology and Management 2021, 486, 118984.

50. Yong-Ju, L.; Go-Eun, P.; Hae-In, L.; Chang-Bae, L. Stand age-driven tree size variation and stand type regulate aboveground biomass in alpine-subalpine forests, South Korea. Science of The Total Environment 2024, 915, 170063, ISSN 0048-9697, 10.1016/j.scitotenv.2024.170063.

51. Hlásny, T.; Barka, I.; Kulla, L.; Bucha, T.; Sedmák, R.; Trombik, J. Sustainability of forest management in a Central European mountain forest: the role of climate change. Reg Environ Change 2015, doi:10.1007/s10113-015-0894-y

52. Dymond, C. C.; Beukema, S.; Nitschke, C. R.; Coates, K. D.; and Scheller, R. M. Carbon sequestration in managed temperate coniferous forests under climate change. Biogeosciences 2016, 13, 1933–1947, 10.5194/bg-13-1933-2016.

53. Pretzsch, H.; del Río, M.; Arcangeli, C. et al. Forest growth in Europe shows diverging large regional trends. Sci Rep. 2023, 13, 15373. 10.1038/s41598-023-41077-6

54. Krejza, J.; Cienciala, E.; Světlík, J.; Bellan, M.; Noyer, E.; Horáček, P.;…Marek, M. V. Evidence of climate-induced stress of Norway spruce along elevation gradient preceding the current dieback in Central Europe. Trees 2021, 35, 103–119.

55. Del Castillo, M. E.; Zang, C.S.; Buras, A. et al. Climate-change-driven growth decline of European beech forests. Commun Biol. 2022, 5, 163. 10.1038/s42003-022-03107-3

56. Gong, X.; Yuan, D.; Zhu, L. et al. Long-term changes in radial growth of seven tree species in the mixed broadleaf-Korean pine forest in Northeast China: Are deciduous trees favored by climate change?. J. For. Res. 2024, 35, 70. 10.1007/s11676-024-01725-7

57. Bennett, A. C.; McDowell, N. G.; Allen, C. D.; & Anderson-Teixeira, K. J. Larger trees suffer most during drought in forests worldwide. Nature plants 2015, 1(10), 1–5.

58. Edward, H. H.; Michael, M.; Trisha, I. H.; Michael, E. U. Recent climatic drying leads to age-independent growth reductions of white spruce stands in western Canada. Glob Change Biol. 2017,23:5297–5308. 10.1111/gcb.13795

59. Reyer C.P.O. Forest Productivity Under Environmental Change—a Review of Stand-Scale Modeling Studies. Current Forestry Reports 2015, 1, 53–68, 10.1007/s40725-015-0009-5

60. Terrer, C.; Vicca, S.; Hungate, B.; Phillips, R. P.; Prentice, I. C. Mycorrhizal association as a primary control of the CO2 fertiliza-tion effect. Science 2016, 353(6294), 72–74. 10.1126/science.aaf4610

